# Structural Basis for Retron Co-option of Anti-phage ATPase-nuclease

**DOI:** 10.1101/2025.05.10.653283

**Authors:** Bing Wang, Renee Hoffman, Ya-Ming Hou, Hong Li

## Abstract

Retrons are recently identified bacterial defense systems that employ a tripartite of reverse transcriptase, non-coding RNA (ncRNA) and its derived multi-copy single stranded DNA (msDNA) to sequester effector activity. Phage invasion activates retrons, triggering effector activity and inducing abortive infection and cell growth arrest. Ec78 distinguishes itself from other retrons by leveraging the Septu defense system, a stand-alone ATPase-nuclease pair (PtuAB), by reshaping its phage sensing and molecular assembly processes. To elucidate the evolutionary adaptation of PtuAB by Ec78, we determined electron cryomicroscopy structures of Ec78 as well as the retron-displaced PtuAB at near atomic resolutions. We show that the Ec78-associated ATPase, PtuA, acquired unique elements that enable its interactions with the reverse transcriptase and the msDNA, as well as self-assembly when displaced by the retron. Supported by biochemical and mutational analysis, we also show that the retron-displaced PtuAB forms a tetramer, unlike its stand-alone counterpart, that indiscriminately restricts the host but, in the presence of the retron, confers a well-controlled immune response, eliciting ATP hydrolysis- and msDNA-regulated targeting to host tRNA^Tyr^. Our studies reveal an evolutionary principle for retrons to co-opt conserved enzyme modules for defense in response to different cellular needs.

**Highlights:**

- Ec78 reverse transcriptase and msDNA sequester PtuAB toxicity
- Ec78 PtuAB heterocomplex targets host tRNA^Tyr^
- ATPase and HNH activities are required for Ec78 PtuAB function
- Ec78 PtuA evolved novel domains for binding retron and for self-assembly
- Ec78 requires distinct ncRNA and msDNA processing for binding PtuAB
- Ec78 retron facilities efficient human genome editing

## Introduction

Once considered selfish genetic elements,^1^ the widely distributed bacterial genetic elements known as retrons have recently been shown to play a role in anti-phage defense.^2, 3^. Retrons protect the bacterial hosts from invading phages through a broad spectrum of effector proteins that include membrane proteins, DNA-binding proteins, metabolic enzymes, ATPases and nucleases.^3^ Investigating how retrons and their effectors contribute to this collaborative process under different selection pressures would reveal their evolutionary capacity and better serve the retron-based biotechnology applications.

Retrons encode a reverse transcriptase (RT) and a non-coding RNA (ncRNA, also referred to as msrRNA) that serves both as the template and the primer for the synthesis of the a multi-copy single-stranded DNA (msDNA, also referred to as RT-DNA) by the RT, leading to expression of the RT, the ncRNA and the msDNA that form the tripartite system^4^. The msDNA is covalently linked to the initiation guanine nucleotide downstream of the conserved UUA motif of the ncRNA via a 2′-5′ linkage (branched). In some cases, the msDNA is separated from the ncRNA by the host XseA-XseB exonuclease (ExoVII) prior to the retron-mediated defense^4, 5^. Furthermore, the template region of the ncRNA is processed by the host RNase H, a process required for the retron-mediated defense^6^, leaving behind a short template:primer duplex with the msDNA. The mature RT-ncRNA-msDNA tripartite sequesters and neutralizes the cytotoxic activities of the effectors that are often encoded as polycistrons along with retrons themselves ^3, 4, 7^. Upon being activated by phage infections, the effector activities lead to abortive infection and cell death. Like CRISPR-Cas, the retron systems provide bacteria with another non-coding RNA-mediated defense against invading phages^8^. The unique production of msDNA by retrons has been harnessed for genome engineering as well as prime-editing, in conjunction with CRISPR-Cas9, across diverse systems.^9, 10^

Retron Ec78 (retron-Eco7) is a Type I-A retron. Of the 11 known types of retrons, Type I-A retrons are unique in partnering with the stand-alone Septu defense system that is an aggregate of PtuA and PtuB proteins (collectively PtuAB) ^2, 3, 11^. The recruitment of PtuAB by Ec78 or other Type I-A retrons suggests that the Septu- and the Ec78-mediated defense are regulated differently. Indeed, unlike Septu that is sensitized by the phage tail fiber^12^, Ec78 produces msDNA, which is consistent with its response to DNA targeting proteins^13^. Despite this difference, the Septu- and the Ec78-associated PtuAB share a high degree of similarity. Both PtuA proteins belong to the family of ABC ATPases^14^ whereas both PtuB proteins share a conserved HNH domain characteristic of the ββα-Me (metal) superfamily of nucleases^15^. The ABC ATPase domain links PtuA to the overcoming lysogenization defect (OLD) nucleases that also function in anti-phage defense and the Mr11-Rad50 proteins for DNA repair^14^. Like a recently characterized OLD nuclease complex, GajA-GajB, whose nuclease function is inhibited by ATP^16, 17^, the stand-alone PtuAB also shows reduced DNA nicking activity as ATP level increases^18^. However, the dependence of Ec78-associated PtuA on ATP remains unknown and its PtuAB targets host tRNA^Tyr^ ^19^. The distinct hosts substrates may explain in part why the stand-alone PtuAB targets a broad spectrum of phages^11, 12^ whereas the Ec78-associated PtuAB is specific for the T5 phage^12^. The molecular basis underlying the defense function of the Ec78 retron remains unclear. Importantly, the evolutionary process by which PtuAB was co-opted into the more complex retron system under the control of the RT, the ncRNA and the msDNA tripartite assembly is elusive.

To elucidate the evolutionary basis for retrons to acquire the stand-alone defense systems, we performed electron cryomicroscopy (cryoEM) and biochemical analysis on the *E.coli* ECONIH5 retron Ec78 (**Fig. 1a & 1b**). Ec78 encodes an unprocessed ncRNA of 144 nucleotides (nts) in length, including 8-nt flanking inverted repeats (IRa1 and IRa2) and four RNA stem loops (stem loops I-IV) of varying sizes. The largest stem loop, stem loop IV, with a 22 base-paired stem, serves as the primary template for msDNA synthesis (**Fig. 1a**). Following msDNA synthesis, stem loop IV is removed by RNase H, leaving behind the ncRNA without stem loop IV, a short template:primer duplex, and the msDNA (**Fig. 1a**). We showed that the Ec78 ncRNA is further processed on retron, leaving behind 40 nts without stem loop I and the two inverted repeats. In addition, the msDNA is debranched, which enables PtuAB to bind. We also showed that, unlike the stand-alone PtuAB, the PtuAB heterocomplex is the cytotoxic unit of the Ec78 retron requiring both the ATPase and HNH activities. We identified the structural elements acquired by PtuAB that enables its switch from the stand-alone to the retron-based defense, thereby illuminating the adaptative process for the Ec78 retron.

**Fig. 1.**
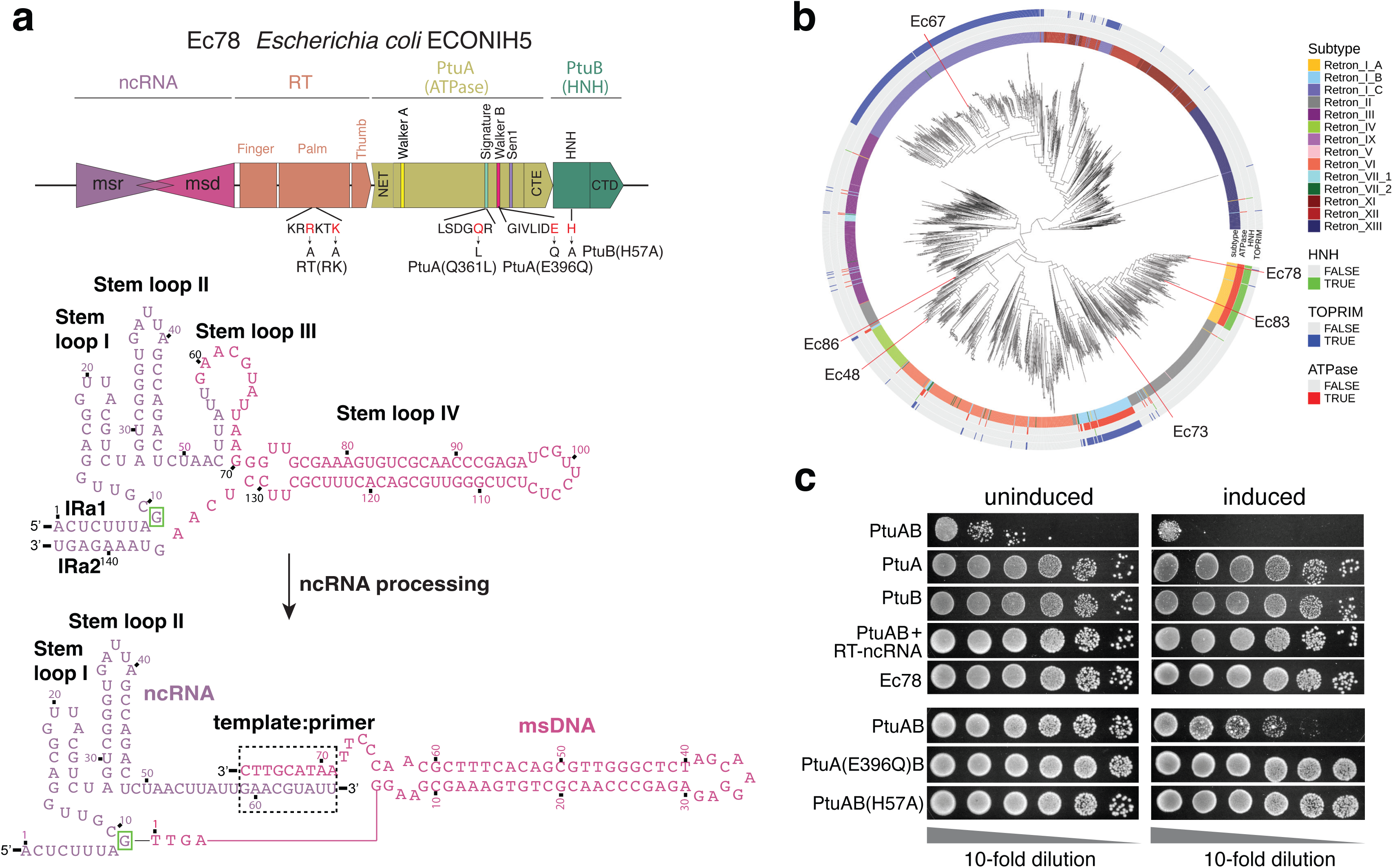
Schematic of Ec78 retron coding sequence features and defense activities. **a**. key features of the Ec78 retron components. Upper, amino acid residues of the motifs within the reverse transcriptase (RT), PtuA, and PtuB relevant to the study are listed. Mutated residues are colored in red. Lower, secondary structures of the non-coding RNA (ncRNA) and msDNA before and after processing. Key regions are colored and labeled. **b**. Co-phylogenetic analysis of retron RTs and their effectors. Retron types are color-coded and labeled. The effectors containing the HNH, TOPRIM or ATPase domains are color-coded and labeled. Some known retrons are marked. **c**. Results of cell growth analysis in DH10B cells transformed with plasmids encoding various Ec78 components and variants of PtuAB. “induced” indicates the plates containing IPTG; “Ec78” denotes the transformation of the single plasmid encoding all retron-effector components; “PtuAB+RT-ncRNA” denotes co-transformation of the plasmids encoding RT and ncRNA, respectively. The top five and bottom three rows are two independent experiments and thus contain respective PtuAB transformation controls.

## Results

### PtuAB complex is the retron-sequestered toxin unit

It was previously shown that the PtuAB components of Ec78 induce cell growth arrest^19^. To understand how each of the PtuAB enzymatic functions contributes to this activity, we conducted growth assays on cells expressing different combination of Ec78 components and their ATPase and HNH deficient variants. We constructed expression plasmids encoding the intact retron, RT-ncRNA, PtuAB, PtuA, PtuB, and their ATPase or HNH deficient variants respectively. Individual plasmids were transformed into the DH10B cells and selected under appropriate antibiotics (**Fig. 1c**). Consistently, expression of PtuAB, but not the intact retron or PtuA and PtuB individually, in DH10B cells severely impaired growth, indicating that PtuAB is the toxin unit (**Fig. 1c**). Furthermore, mutation of the Walker B motif residue Glu396 (E396Q) of PtuA or the catalytic residue His57 of the HNH domain (H57A) of PtuB in PtuAB co-expression cells rescued the growth defects (**Fig. 1c**), suggesting that both the ATPase activity and the nuclease activity are required for cell growth arrest. These results confirm PtuAB as the functional defense unit and both the ATPase and HNH nuclease activities are necessary for the function, a conclusion that is in stark contrast to that of the stand-alone PtuAB system whose activity is strictly controlled prior to phage invation^18, 20^. Therefore, in the Ec78 retron, the RT-ncRNA-msDNA entity exerts control over the adapted PtuAB for its defense function. Upon phage infection, the PtuAB displaced by Ec78 indiscriminately impairs host targets, leading to abortive immunity.

### The PtuAB complex targets host tRNA^Tyr^

Previous efforts to identify phage determinants that evade Ec78 defense identified tRNA^Tyr^ as a potential target ^19^. To confirm that PtuAB targets host tRNA^Tyr^ specifically, we co-transformed plasmids that overexpress tRNA^Tyr^-GTA-1, tRNA^Tyr^-GTA-2, tRNA^Ala^, tRNA^His^, tRNA^ser^, tRNA^Arg^, tRNA^Met^, or no tRNA (null) along with that expressing PtuAB into BL21(DE3) cells to increase the expression levels (**Fig. 2a**). We showed that overexpression of both tRNA^Tyr^ (GTA-1 and GTA-2) molecules, but not the null or other tRNAs, notably reduced PtuAB toxicity, indicating PtuAB specifically targets tRNA^Tyr^ (**Fig. 2a & Suppl. Fig. 1a**), consistent with the previous finding that mutant phages overexpressing tRNA^Tyr^ evade retron Ec78 ^19^. The toxicity of PtuAB in the non-expression cells DH10B can also be rescued by the expression of tRNA^Tyr^ and tRNA^Ala^ to a lesser degree (**Suppl. Fig. 1b**).

**Fig. 2.**
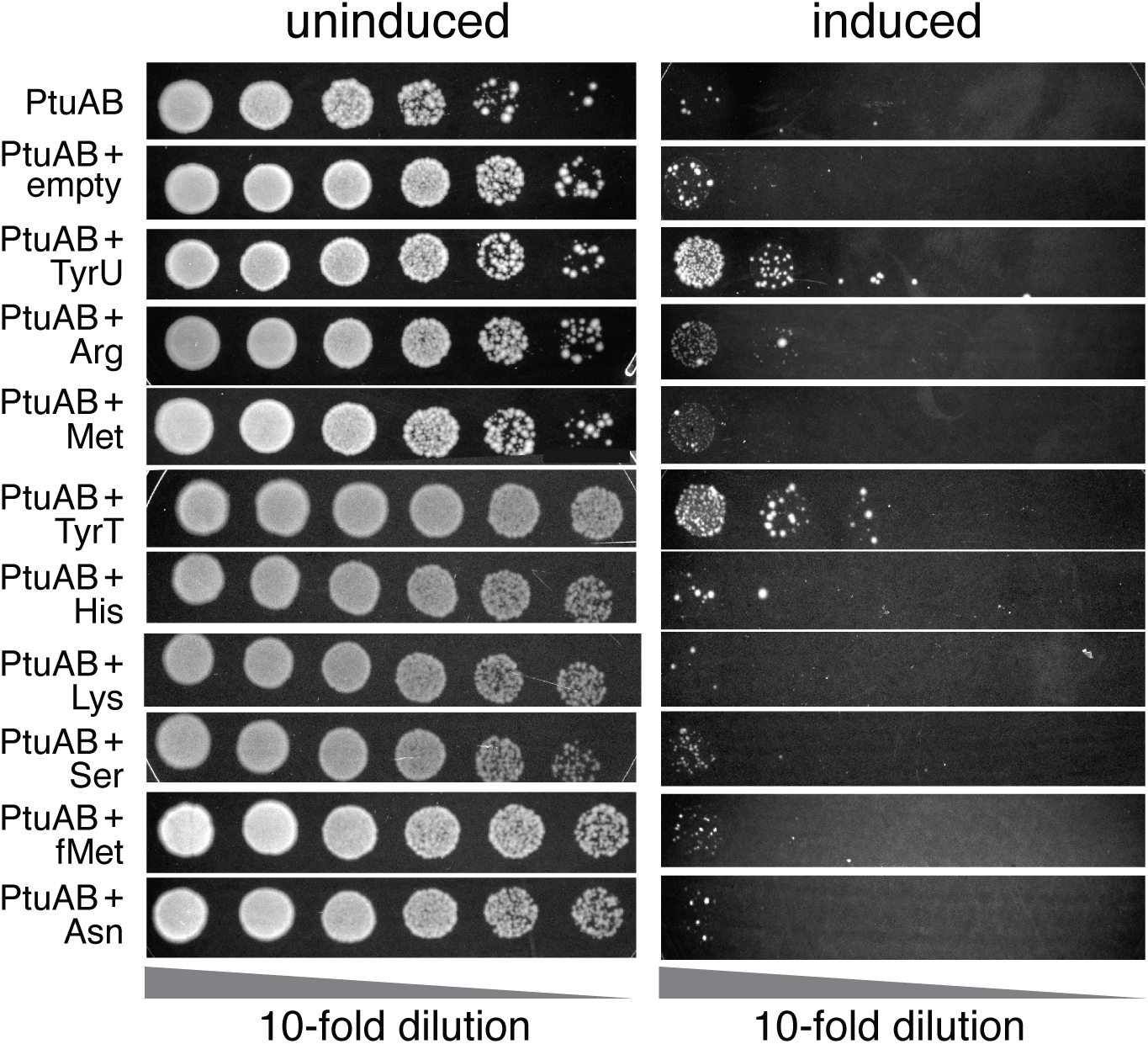
Cell growth assays reveal Ec78 targets host tRNA^Tyr^. Results of cell growth analysis in BL21(DE3)pLysS cells co-transformed with the plasmid encoding PtuAB and that encoding various E. coli tRNAs. “induced” indicates the plates containing IPTG.

We further reconstituted the wild-type PtuAB complex by co-purifying them from the cells overexpressing tRNA^Tyr^ and the intact Ec78 complex including their catalytic variants and verified that the wild-type PtuA or its complexes are active in ATP hydrolysis (**Suppl. Fig. 2**). We subsequently performed *in vitro* tRNA cleavage assays. To our surprise, neither PtuAB nor PtuB or the DNase I-treated Ec78 was able to degrade in vitro transcribed tRNA^Tyr^-GTA-2 (**Suppl. Fig. 1c**). Furthermore, the DNase I-treated Ec78 also did not degrade the native tRNA^Tyr^ isolated from E coli^21^ in the presence or absence of divalent ions or ATP (**Suppl. Fig. 1d**). It is possible that an unidentified host factor is further required for the observed targeting activity in cells.

The tRNA targeting activity by Ec78 differs from the stand alone Septu system that has been reported to target DNA^18, 20^. The Ec78 PtuAB tRNA targeting activity is more consistent with that of the ATPase-TOPRIM nuclease pair, AriA-AriB, of the phage antirestriction induced system (PARIS) system, which is also a retron-independent abortive immunity system^22, 23^. However, AriB contains the TOPRIM-like domain and functions when released from AriA^22, 23^ whereas PtuB contains a HNH domain and requires PtuA for function.

### Molecular basis for Ec78 retron to sequester PtuAB

To unveil the molecular basis for Ec78 to sequester PtuAB, we carried out cryoEM structural analysis of the purified retron-effector complex. The. sample preparation process indicated a sensitivity of assembly to ATP hydrolysis. Whereas the wild-type retron displayed significant heterogeneity in assembly (**Suppl. Fig. 2a**), especially due to PtuB displacement, that containing the ATPase deficient mutant, E396Q in PtuA showed reduced heterogeneity and allowed its structural determination (**Suppl. Fig. 3 & Suppl. Table 1**).

The cryo-EM map revealed well-resolved density for both the ncRNA and the msDNA, enabling unambiguous assignment of their nucleotides (**Fig. 3 & Suppl. Fig. 3**). Surprisingly, only stem loop II (rU27 to rU48), a linker (rC49-rU58) and the template region (rG59-rU67) base paired with msDNA (dA69-dC77) of the ncRNA are observed. Furthermore, the msDNA is debranched at its 2′-5′ linkage to the initiation guanine nucleotide and lacks the first four nucleotides (dT1-dA4) (**Fig. 3a**), unlike that previously observed in the Ec86 retron^24, 25^. We confirmed the processing of ncRNA and msDNA by analyzing the nucleic acids co-purified with either the intact retron or RT alone by examining the nucleic acids from the cell extracts or co-purified with proteins on PAGE gels (**Fig. 3b**). We further showed that the processing of both nucleic acids does not require the presence of PtuAB as the processed products are also present in the RT alone sample (**Fig. 3b**). To assess if RT influences the processing activities, we mutated Lys143 and Arg146 of the RT (RT(RK)) that are found near the msDNA cleavage site. We showed that the RT(RK) variant reduced processing (**Fig. 3b**), and its retron assembly no longer sequesters the PtuAB toxicity (**Suppl. Fig. 4**), suggesting that, though host enzymes are likely responsible for the observed processing activities, RT exerts a notable influence on this process.

**Fig. 3.**
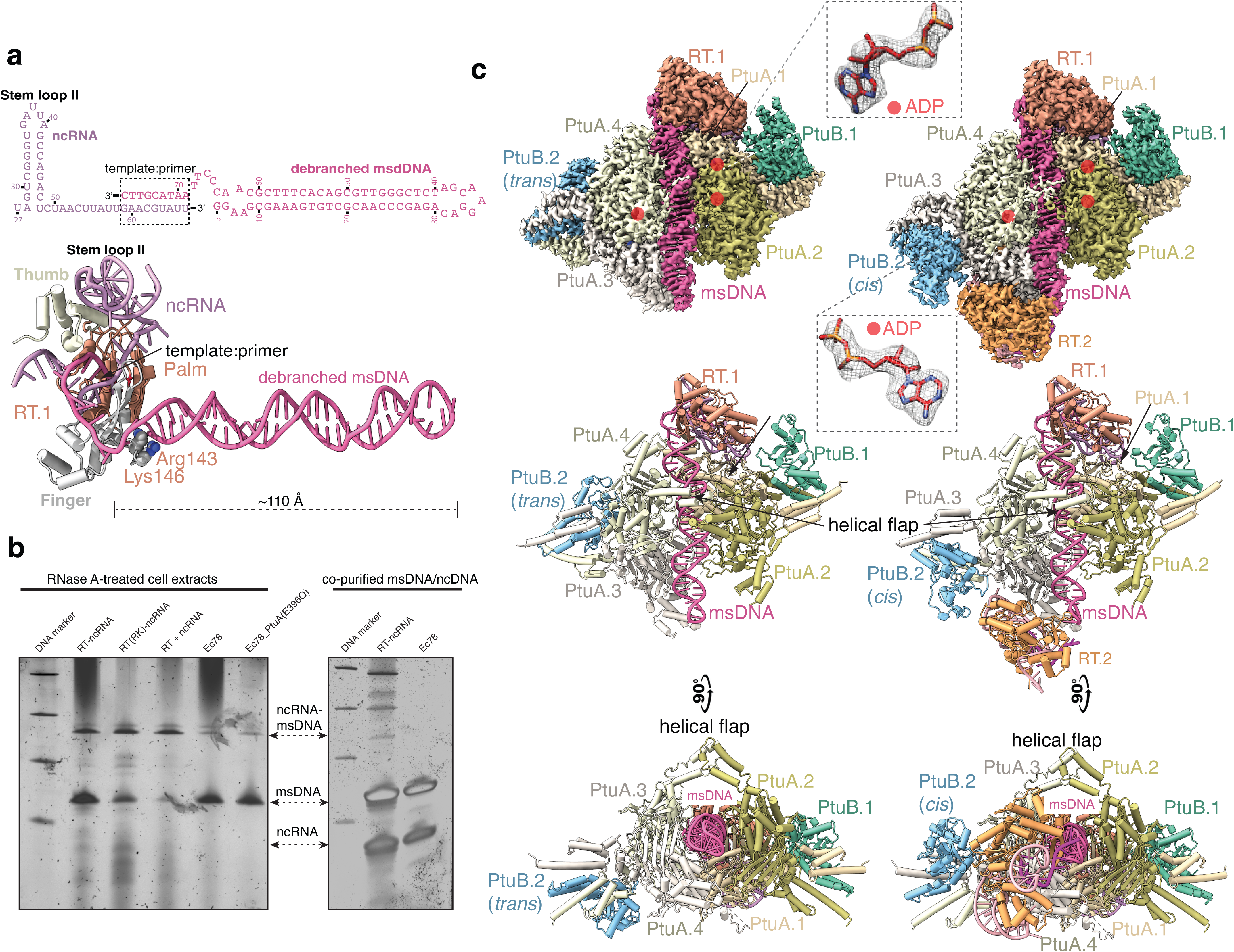
Structure overview of the Ec78 retron and analysis on ncRNA and msDNA processing. **a**. The secondary and tertiary structures of the ncRNA and msDNA observed in the Ec78 structures. Only the reverse transcriptase (RT) is retained for simplicity with its characteristic domains labeled. **b.** Gel analysis results of the nucleic acids extracted from the purified Ec78, RT-ncRNA, and RT(R143A/K146A)-ncRNA (RT(RK)) (right) or from the RNase A-treated cell extract (left). **c**. Density maps (upper) and cartoon representations (lower) of the Ec78 retron complex. The 4:2:1 complex is shown to the left with one PtuB in trans with respect to the top RT and the 4:2:2 complex is shown to the right with the corresponding PtuB in cis with respect to the top RT. Orange dots indicate the bound ADP molecules. Insets show the close-up views of the density around the two selected ADP molecules in the 4:2:1 complex.

The retron-effector particles are classified into two major classes. The first is a 4:2:1 ratio of PtuA, PtuB, and RT-ncRNA-msDNA, with four PtuA subunits, two PtuB subunits, and one RT-ncRNA-msDNA complex (**Fig. 3c**). The second class exhibits a 4:2:2 stoichiometry, featuring an additional RT-ncRNA complex at the distal end of the msDNA hairpin, where only the terminal 12 msDNA nucleotides (dC65-dC77) are visible (**Fig. 3c**). This suggests that the complete the Ec78 retron assembly accommodates a single msDNA molecule. There are other classes with either reduced PtuB occupancy or different combinations of PtuB orientations (cis vs. trans to RT) without other structural differences. These classes led to reconstructions at lower resolutions and are thus not modeled (**Suppl. Fig. 3 & Suppl. Table 1**). Interestingly, though we used the E396Q PtuA variant in structural determination, which showed significantly reduced ATPase activity (**Suppl. Fig. 2c**), ADP instead of ATP was trapped in three of the four subunits (**Fig. 3c**). This result contrasts with the stand-alone PtuAB structures in which ATP molecules are trapped, consistent with its lack of ATP hydrolysis ^18, 20^. Since there are no other differences in molecular interactions among these classes of structures, we focus the subsequent structural analysis primarily on the 4:2:1 assembly.

A single RT subunit forms the head of the Ec78 retron assembly from which a long msDNA hairpin emerges to form the central spine of the assembly (**Fig. 3c**), unlike the previously observed Ec86 retron structure that is capped by RT dimers and bordered by two antiparallel msDNA^24, 25^. In the 4:2:2 class, a second RT is positioned at the distal end of the msDNA hairpin but with its own msDNA largely disordered, leading to an essentially same architecture (**Fig. 3c**). The lack of the dimerization α-helix at the N terminus of Ec78 RT is responsible for the different oligomerization state than that of Ec86.

The 3’-overhang of the msDNA is anchored to the RT-bound ncRNA via a short RNA:DNA duplex (template:primer) that feeds into the catalytic Palm domain of RT while its hairpin extends approximately 110 Å from the RT-ncRNA head (**Fig. 3a**). Four PtuA subunits, forming a dimer of dimers, construct a diamond-shaped platform with a central groove that adheres along the length of the msDNA hairpin, while two PtuB proteins cap the outer edges (**Fig. 3c**). Should the IRa1 and stem loop I of the ncRNA and the 2′-5′ linkage of the msDNA is retained, they would obstruct the assembly of PtuAB, suggesting that removal of these ncRNA elements, including msDNA debranching, is a required mechanism for Ec78 to achieve sequestration of the toxicity.

The PtuA platform forms an extensive interface with the msDNA, totaling ∼2000 Å^2^ buried solvent accessible area (**Fig. 4a**). However, the contacts are non-specific and mostly occur between the loops in the helix-grip fold of PtuA and the minor groove of the msDNA (**Fig. 4b & 4c**). Most of these amino acids are positively charged and are not conserved in Septu PtuA (**Suppl. Fig. 5a**). In addition, a unique helix-loop-helix insertion to the RecA-like nucleotide binding domain of PtuA forms a helical flap with the corresponding element of an opposing PtuA subunit that encircles the msDNA (**Fig. 3c** & **Fig. 4b**). Thus, the assembly of PtuA with msDNA is based on shape complementarity, similar to other ABC ATPases, such as the Rad50 ATPase^26^, that act on DNA nonspecifically. Importantly, loop forming a number of contacts with msDNA (m loop) is significantly longer in the stand-alone PtuA, which would obstruct its assembly with DNA (**Fig. 4b & Suppl. Fig. 5a**). In addition, the helical flap encircling the msDNA is deleted in the stand-alone PtuA (**Fig. 4b & Suppl. Fig. 5a**). The distinct elements in Ec78 PtuA evolved for binding msDNA enables it to partner with Ec78, reflecting a unique evolutionary path for retron acquisition.

**Fig. 4.**
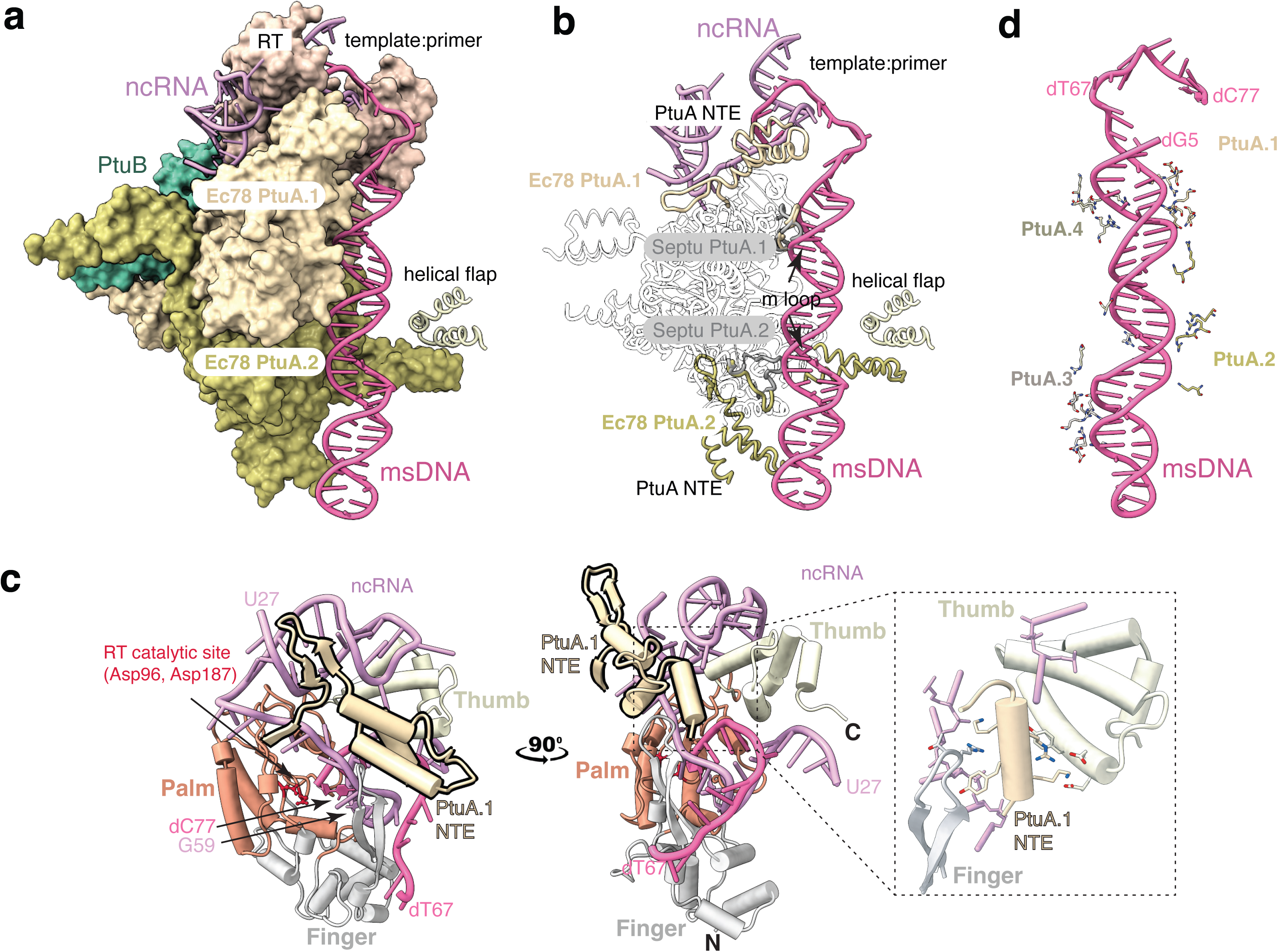
Interactions between PtuA and Ec78 components in Ec78 structures. **a.** Binding of msDNA to the central groove formed by PtuA. RT, PtuA, and PtuB molecules are shown in surface representation. The other dimer of PtuAB is omitted for clarity. **b**. Superimposition of the Ec78 PtuA to the Septu PtuA (gray ribbons) structures reveal unique protein elements in Ec78 PtuA evolved for contacting msDNA (m loop and helical flap) and the RT-ncRNA complex (N-terminal extension, NTE). The loop equivalent to the m loop in Septu PtuA would obstruct its ability to contact msDNA in the same manner. **c**. PtuA residues, mostly positively charged, making close contacts (within 3.5 Å) with the msDNA. These residues are indicated in Supplementary Figure 5. **d.** Close-ups in two orthogonal views illustrating structural features of PtuA interactions with the RT-ncRNA-msDNA tripartite. The characteristic domains of the RT are indicated and color coded. Nucleotides of the ncRNA and msDNA are labeled. NTE denotes the N-terminal extension of PtuA. Inset highlights the specific contacts of first α-helix of NTE with both the Finger and Thumb domains.

The association of PtuA with Ec78 retron is further enhanced by its contacts with RT, which buries ∼690 Å^2^ solvent accessible surface (**Fig. 4c**). This interaction is largely mediated by the N-terminal extension (NTE) of PtuA, an element also distinctly absent from the stand-alone PtuA (**Suppl. Fig. 5a**). The first α-helix of NTE is wedged between the characteristic Finger and the Thumb domains of RT, forming several specific protein-protein contacts (**Fig. 4c**). In addition, the first α-helix of the NTE also contacts the ncRNA (**Fig. 4c**), though, non-specifically, which further stabilizes the PtuA-retron association. The observed NTE-RT-ncRNA interaction is another significant feature acquired by the Ec78 PtuA.

### Mechanism of PtuAB activation

To determine if there are distinct sequence features in the PtuAB proteins adapted to function with Type I-A retrons, we performed phylogenetic analysis of both proteins. The HNH nuclease PtuB revealed a somewhat scattered distribution between retron-associated and retron-independent sequences (**Fig. 5a**), suggesting minimal evolutionary pressure for co-opting stand-alone PtuB to function within the Type I-A retrons. In the retron structures, PtuB employs its helical C-terminal domain (CTD) to interact with the C-terminal extension of PtuA (**Fig. 5b**). A pair of CTEs from each PtuA dimer sandwiches one PtuB subunit, forming a shared nine-helix bundle where three helices are from PtuB (**Fig. 5b**) Both CTD of PtuB and CTE of PtuA seem to be well conserved between the Ec78 and Septu systems (**Suppl Fig. 5**). Furthermore, a comparison of the overall structure (**Fig. 5c**) and the active site (**Fig. 5d**) of Ec78 retron PtuB with stand-alone PtuB and the HNH domain of AceCas9 ^27^ indicates a high degree of structural similarity, which, however, seems to contradict to their different nucleic acid targets.

**Fig. 5.**
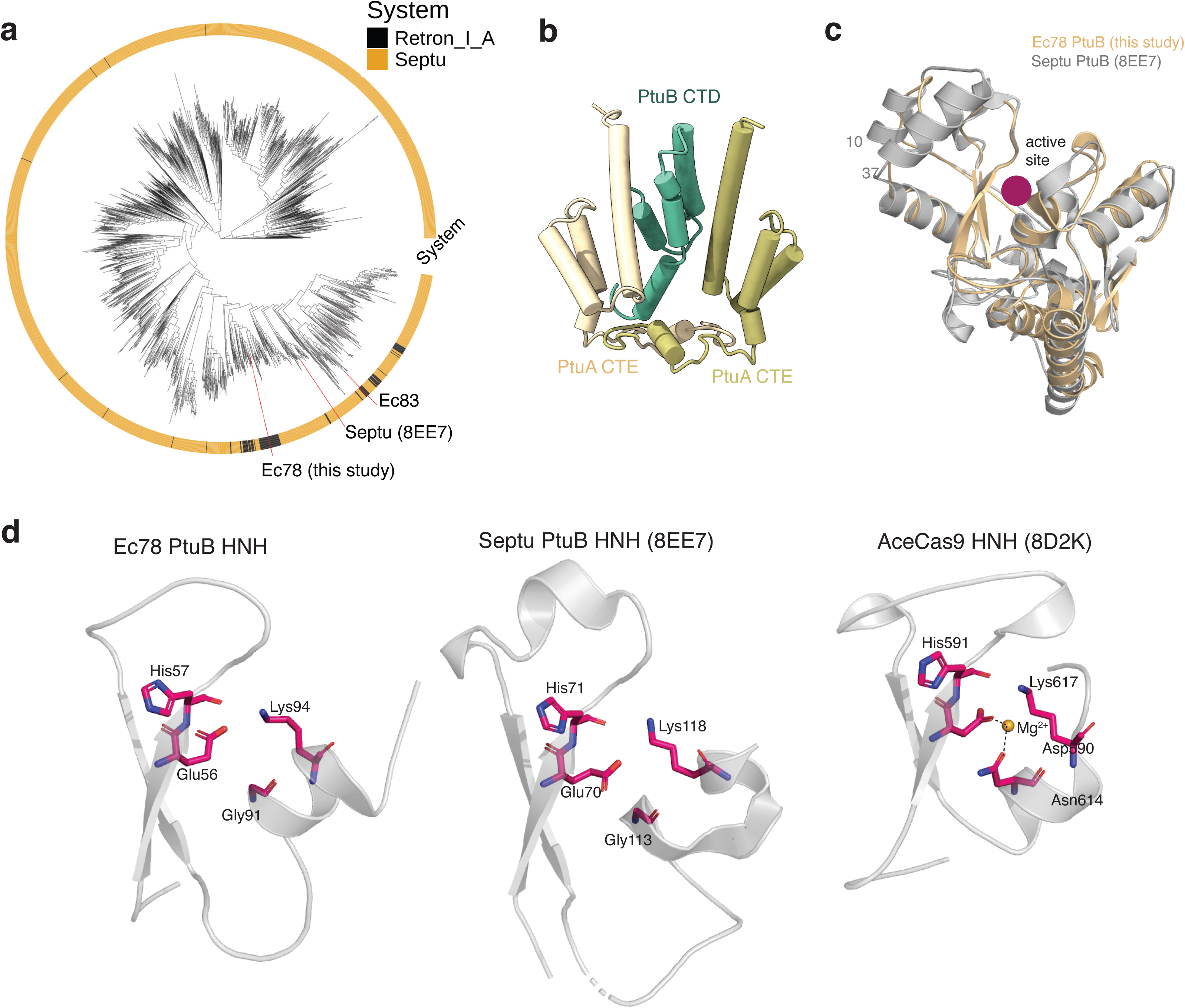
Phylogenetic and structural analysis of PtuB. **a.** Phylogenetic analysis of both retron Type I-A (black) and Septu (orange) PtuB sequences. **b**. Close-up view of the C-terminal domain of PtuB (CTD) interacting with a pair C-terminal extension (CTE) of PtuA. **c.** Superimposed PtuB structures between the Ec78 (beige) and the Septu (gray, PDB code: 8EE7) systems. The catalytic site is marked by a red dot. The region displaying the largest difference is marked by Ec78 PtuB residues. **d.** Comparison of the HNH domain active site among Ec78 PtuB, Septu PtuB (PDB code: 8EE7) and the Cas9 from *Acidothermus cellulolyticus* (PDB code: 8D2K).

Unlike PtuB, the ATPase PtuA exhibits distinct clustering between the retron-associated and retron-independent sequences (**Fig. 6b**), suggesting that the evolution of the PtuA subunit from the stand-alone Septu system facilitated its integration into retron function. This is further supported by the observed structural features unique to the retron-associated PtuA for binding the RT-ncRNA-msDNA tripartite complex that are distinctively absent from the stand-alone PtuA (**Fig. 4b & Suppl. Fig. 5a**).

**Fig. 6.**
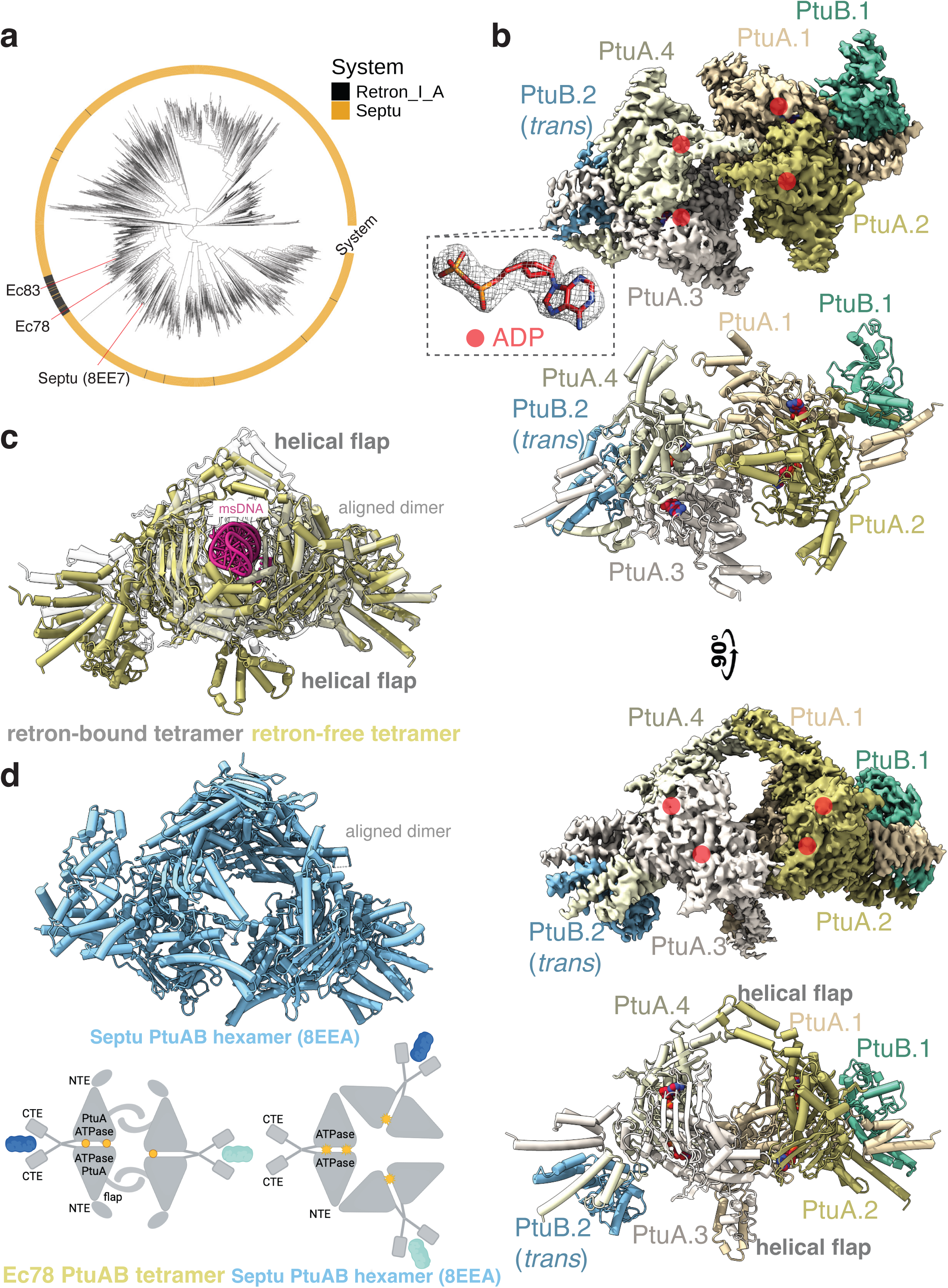
Phylogenetic analysis of PtuA and structural analysis of the retron-free PtuAB. **a.** Phylogenetic analysis of both retron Type I-A (black) and Septu (orange) PtuA sequences. **b.** Density maps and the corresponding cartoon representations of the Ec78 retron-displaced PtuAB complex in two orthogonal views. Orange dots indicate the bound ADP molecules. Inset shows the density map around one of the bound ADP molecules. **c.** Structural comparison between the retron-displaced (orange) and the retron-bound PtuAB (gray), both in tetrameric forms, when a single PtuA subunit is aligned as indicated. The msDNA model is from the 4:2:1 Ec78 structure and shows clashes with the retron-displaced PtuA. **d**. Schematic oligomers of the retron-displaced (tetramer) and the Septu (hexamer) PtuAB. The hexameric structure of Septu PtuAB aligned with a PtuA subunit is shown to the right. Note that PtuB molecules are included in the assembly but not highlighted.

To further analyze the molecular basis for PtuAB activation, we obtained its structure in the absence of the retron. We introduced the Gln361 to leucine mutation within the signature motif (357-LSDGQR-362) of PtuA^14^ (**Suppl. Fig. 5b**) to abrogate the wild-type toxicity, which permitted protein purification and structure determination (**Suppl. Fig. 6 & Suppl. Table 1**). Single particle reconstruction led to a major class of PtuAB particles that was refined to 3.2 Å (**Fig. 6b**).

The PtuAB complex form a similar diamond-shaped platform comprising four PtuA and two PtuB subunits (**Fig. 6b**), significantly different from the hexameric and inflammasome-like assembly of the stand-alone PtuAB^18^. The retron-displaced Ec78 PtuAB assembles largely through dimerization of two helical flaps that otherwise encircle msDNA when assembled with retron (**Fig. 6b**). In comparison with the retron-sequestered PtuAB, the retron-displaced PtuAB displays a substantially narrower central groove and a more compressed helical flap (**Fig. 6c**), both of which would exclude msDNA from binding. Interestingly, the retron-displaced PtuAB bearing the Q361L mutation in the ATPase domain also traps ADP as opposed to ATP on all four sites (**Fig. 6b**), consistent with its detectable ATPase hydrolysis activity (**Suppl. Fig. 2c**). Notably, the stand-alone PtuA contains leucine instead of Gln361 at this position and lacks ATPase activity, indicating that the retron-associated PtuA is adapted to have a strong ATPase activity.

The unique involvement of the helical flap in Ec78 PtuA oligomerization prevents it from forming the same inflammasome-like hexameric assembly observed for the stand-alone PtuAB complex^18^ (**Fig. 6d**). This difference may sufficiently explain the differences in both ATP hydrolysis and defense targets between the two PtuAB systems. It is remarkable that through addition of the elements to the well-conserved ATPase core, the PtuAB proteins are co-opted for sensing different phage determinants and targeting different host substrates.

The conformational closure of the retron-displaced PtuAB suggests a possible mechanism of its activation (**Fig. 7**). In the presence of the msDNA and the associated RT-ncRNA, PtuA is locked in a conformation that prevents rapid displacement of PtuB. In the absence of the msDNA, for instance as a result of the phage attack, and in conjunction with ATP hydrolysis, PtuA activates PtuB for its nuclease activity, leading to growth arrest. This msDNA-mediated sequestration mechanism is consistent with the detected phage escape mutants in the D15 DNase^13^.

**Fig. 7.**
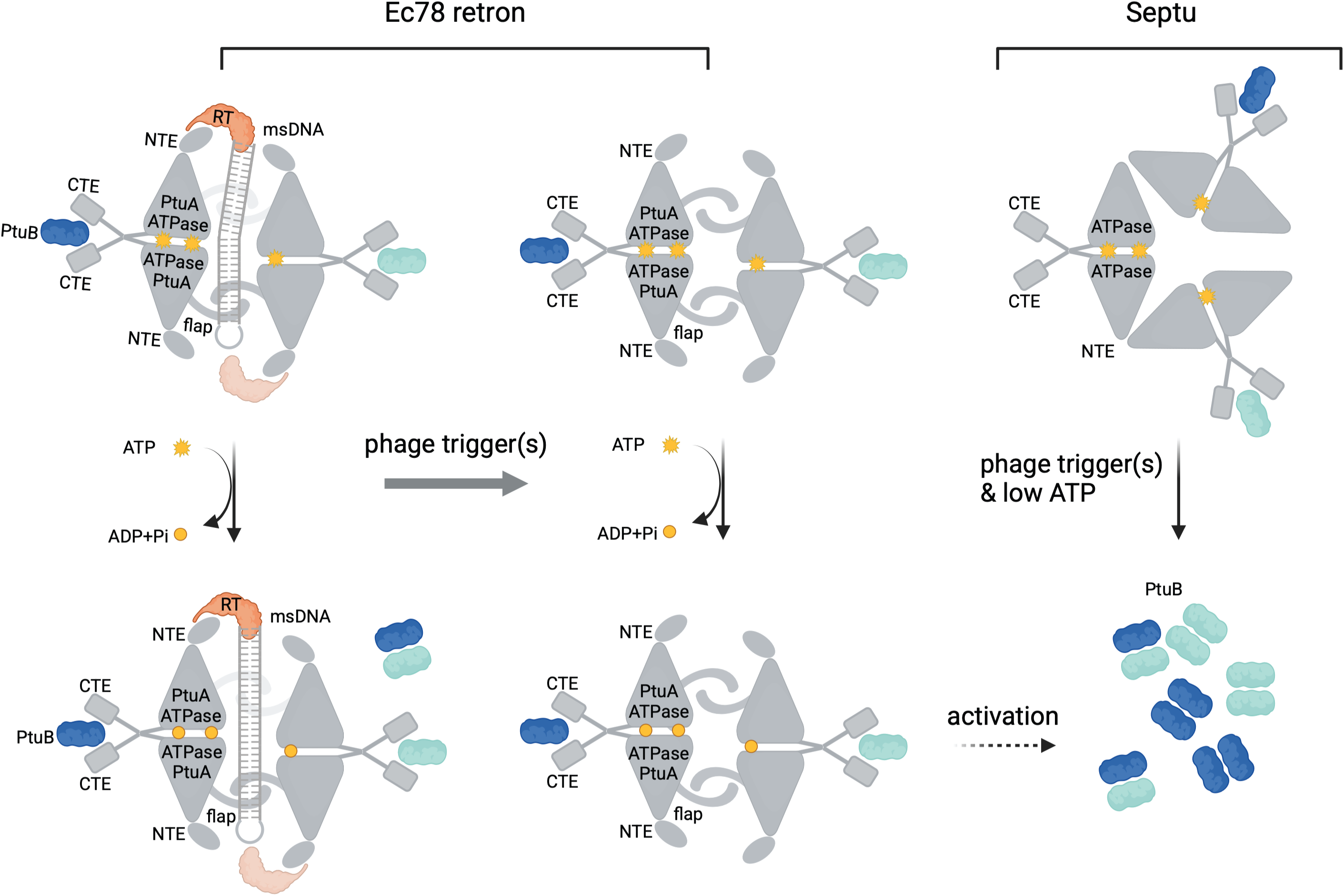
Proposed models of activation for the Ec78 and Septu systems. Left, Ec78 activation model. The RT-ncRNA-msDNA sequestered PtuAB undergoes some release of the PtuB molecules with the help of ATP hydrolysis without causing cell growth arrest. Upon phage infection, the tetrameric PtuAB is released from the tripartite and, in the presence of ATP hydrolysis and a yet to be identified factor, PtuB is released to target tRNA^Tyr^ molecules. Right, Septu activation model. The hexameric PtuAB complex is activated by phage proteins, and upon the reduction of ATP levels, releases PtuB molecules to target genomic DNA.

### Ec78 retron facilities efficient genome editing in human cells

To assess the ability of Ec78 retron for producing DNA to support genome editing in human cells, we transformed Ec78 retron that encodes a msDNA with flanking homology arms to a EMX1 site (**Fig. 8a**) into HEK293T cells that expresses an endogenous SpyCas9 targeting the same EMX1 site. Previous genome editing studies using over 100 retrons found that efficient production of msDNA led to replacement of the target site by the retron-synthesized msDNA. We thus included a highly efficient retron previously identified, Mestre-1531, a Type II-A retron, targeting the same site as the control. Next generation sequencing showed that Ec78 retron produced at least comparable genome editing efficiency as Mestre-153 (**Fig. 8b**). These results make Ec78 retron the only one that is highly effective in genome editing and with 3D structure information.

**Fig. 8.**
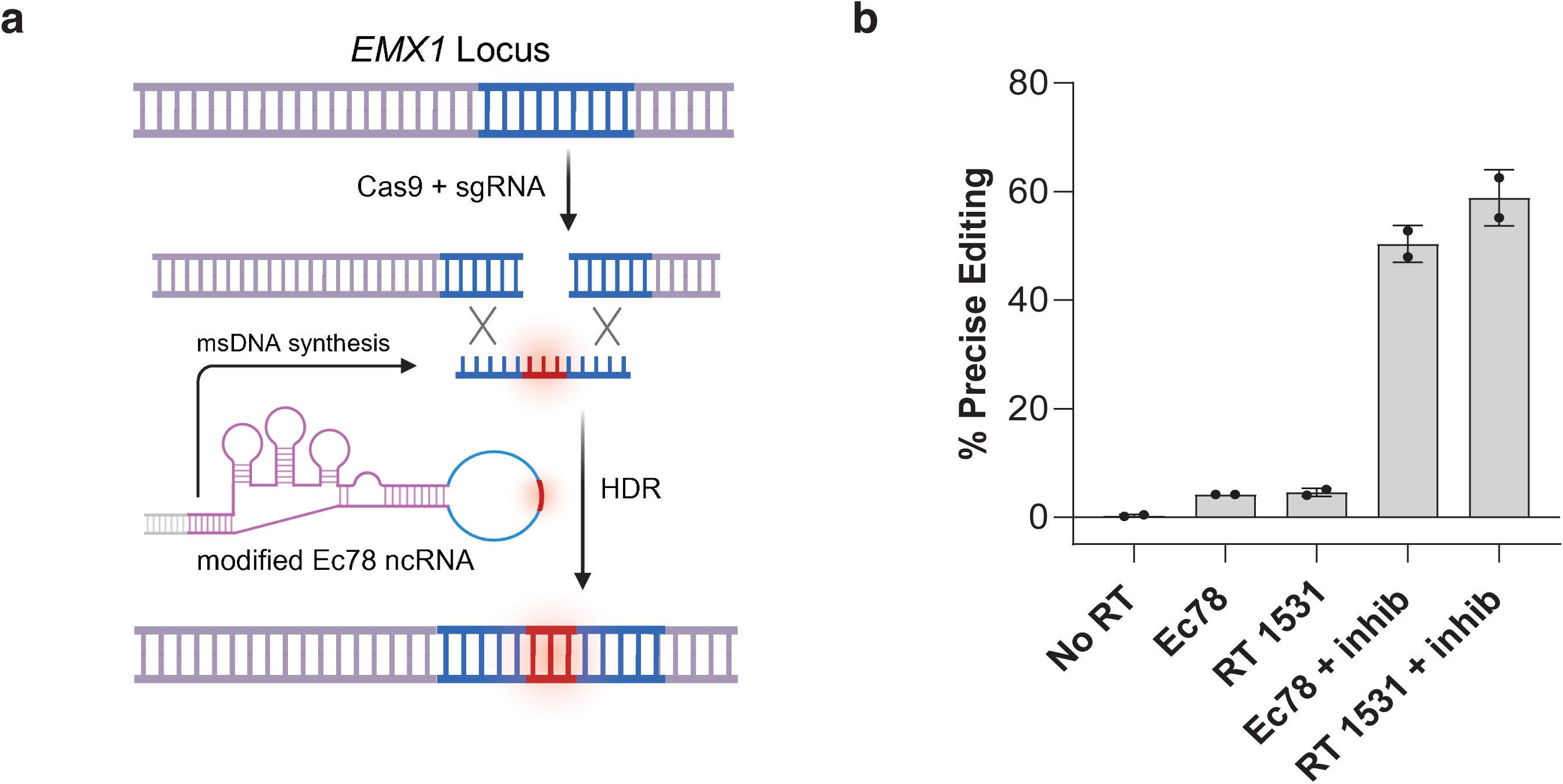
Ec78 retron precise editing. **a.** Schematic of Ec78 retron precise editing targeting an EMX1 site. **b.** Quantification of Ec78 retron precise editing in comparison with no reverse transcriptase control (No RT), in the absence (Ec78) or the presence (Ec78+inhib) of inhibitors, AZD7648 and PolQi2, and the same experiments with the Mestre-1531 retron (RT 1531) with the same inhibitors. Bars represent means ± standard deviations of two biological replicates.

## Discussions

We report structural and functional characterization of the first retron example that co-opts a stand-alone anti-phage system for retron-mediated defense. In contrast to the protein-controlled Septu PtuAB, the Ec78 PtuAB possesses strong ATP hydrolysis activity and is neutralized by the RT-ncRNA-msDNA tripartite. Upon being displaced, the PtuAB indiscriminately restricts the host, leading to growth arrest. The Ec78 PtuAB acquired three protein elements: the N-terminal extension to interact with the RT and ncRNA, a helical flap and a number of positively charged loops to entrap the msDNA. Acquisition of these tripartite interacting elements eliminates its ability to self-assemble into the inflammasome-like complex as its stand-alone counterpart in Septu that allows a protein-mediated control, such as sensing the phage tail fiber. The difference in PtuAB assembly is likely also responsible for the different host targets observed for the Septu PtuAB, which targets genomic DNA, and Ec78 that targets host tRNA^tyr^. Interestingly, another type I-A retron, Ec83, that also partners with PtuAB, does not seem to target host tRNA^19^, raising the possibility that the PtuAB system offers a versatile platform for sensing additional phage triggers and targeting different nucleic acids in response to different environmental pressures.

The function of Ec78 retron RT resembles that of Ec86 retron^24, 25^ and that of the diversity-generating retroelements (DGR)^28^, for which three dimensional structures are now available, in using ncRNA to template DNA synthesis. Different from these latter RTs, however, Ec78 RT has evolved features specific for its partner effector. The Ec78 RT protein adopts the characteristic “right-hand” architecture typical of group II intron family of RTs, comprising three distinct subdomains - Finger, Thumb and Palm – that are all preserved in the three RT enzymes. Structural comparison reveals a high similarity between Ec78 and Ec86, with a root mean square deviation (RMSD) of 1.155 over 121 Cα atoms. However, Ec78 RT exhibits two distinguishing features: the absence of an N-terminal helix preceding the Finger domain and a particularly long, positively charged loop within the Palm domain (Lys141-Lys146). In Ec86 RT, the N-terminal helix facilitates RT dimerization, whereas its absence in Ec78 RT accounts for the monomeric state of RT in the Ec78 retron system. The positively charged loop plays a critical role in stabilizing the 5’ end of the msDNA, with the side chain of Arg143 forming a hydrogen bond with the phosphate group of terminal dG5, while Arg142, Arg143 and Lys146 engage in electrostatic interaction with dG5 and dG6. We found that the R143A/K146A mutation impaired msDNA biogenesis required for retron assembly, leading to cytotoxicity when the mutant retron is expressed in cells.

Consistent with the structural data, phylogenetic analyses also revealed co-evolved features that support the specific assembly. We observed distinct clusters of RTs based on types, with only a minor proportion scattered throughout the tree. For instance, the positively charged loop that impacts msDNA debranching is highly conserved among Type I-A retrons, whereas the N-terminal helix required for dimerization is a common feature in Type II retrons (such as Ec86). Notably, this clustering pattern is more pronounced in larger subtypes with a greater number of members, whereas smaller subtypes, with fewer members, exhibit greater dispersion. These findings suggest that co-evolutionary selection plays a dominant role in retron evolution, while modular exchange occurs to a lesser extent, particularly in smaller subtypes. These findings suggest a potential co- evolutionary relationship between RT and its associated effector in retron system, highlighting structural adaptations that contribute to functional specificity within each retron type.

The unique production of msDNA by retrons has been harnessed for genome applications, in conjunction with CRISPR-Cas9, in bacteria, yeast and human cells^10, 29^. Transfecting the plasmid carrying the RT-msrRNA cassette in targeted cells produces msDNA by the RT, providing an amplifiable single-stranded DNA that act as the homology dependent repair (HDR) template to repair the double-stranded break generated by CRISPR-Cas9. Owing to the continuous template production, the retron-mediated HDR is shown to be more efficient than that mediated by synthetic DNA^10^. In addition, retron RTs have been repurposed for prime-editing, in which the fused nickase CRISPR-Cas9 nicks the DNA while the RT produces the desired edited DNA from the prime editing guide RNA (pegRNA).^29^ Retron reverse transcriptases present a promising, compact alternative for prime editing, achieving efficiencies comparable to the larger M-MLV RT through protein engineering^29^. Our results indeed showed highly efficient genome editing with Ec78 retron. However, the implications of the distinctive biogenesis of Ec78 ncRNA and msDNA in genome editing remain unclear, highlighting a need for further research.

## Materials and Methods

### Cloning, Protein Expression and Purification

For protein expression, the operon sequences of *E.coli* ECONIH5, including its native promoter, coding sequences of ncRNA, RT, PtuA and PtuB, were PCR amplified from the pLG008 plasmid^2^. The PCR product was cloned into a modified polycistronic expression plasmid, pST44^30^, where a C-terminal 3C protease-cleavable His-tag was introduced at C-terminus of PtuB by Gibson assembly. For the co-expression of PtuA and PtuB, the coding sequences of PtuA and PtuB were individually cloned into the first and second multiple cloning sites of a modified pRSF-Duet expression vector, where an N-terminus of His-Sumo tag was introduced on PtuA. For the toxicity assay, the sequences encoding PtuA and PtuB were cloned either separately or together into expression vectors. When cloned separately, each gene was inserted into either the pBAD vector or the pRSF-Duet vector. For co-expression, the ptuA and ptuB genes were cloned together either in the pRSF-Duet vector or as part of a co-operonic construct in the pBAD vector. All the mutations were performed by QuikChange mutagenesis method with Q5 DNA polymerases.

The expression plasmids were transformed into either the Nico21 (DE3) (NEB) or the BL21-CondonPlus (DE3)-RIPL (Agilent) cells. Transformed cells were plated onto LB agar plates containing appropriate antibiotics for selection. A single colony from the plate was used to inoculate 50 ml of LB medium supplemented with the same antibiotics and the primary culture was grown overnight at 37 °C. Eight milliliters of the overnight culture were transferred into 1 L of LB medium with a total volume of 6 L prepared for subsequent purification. The cells were grown at 37 °C until the OD_600nm_ reached 1.0. The temperature was lower to 19 °C, and after 30 minutes, the culture was induced with 0.2 mM isopropyl β-d-1-thiogalactopyranoside (IPTG). Induction was allowed to proceed overnight. After overnight induction, cells were pelleted by centrifugation and snap-frozen in liquid nitrogen and stored at −80 °C if not used for purification immediately.

For protein purification of the retron complex, the cell pellets were resuspended and lysed by sonication in 80 ml of lysis buffer (50 mM Tris-Cl, 500 mM NaCl, 5% glycerol, 10 mM Imidazole, 5 mM TCEP). The lysate was clarified by centrifugation, and the supernatant was loaded into a 5 ml HisTrap HP column (Cytiva) using a peristaltic pump. Subsequent washing and elution steps were also performed with the peristaltic pump. The column was sequentially washed with 200 ml of low-imidazole wash buffer (50 mM Tris-Cl, 500 mM NaCl, 5% glycerol, 25 mM Imidazole) followed by50 ml of high-imidazole wash buffer (50 mM Tris-Cl, 500 mM NaCl, 5% glycerol, 40 mM Imidazole). Protein was then eluted with 20 ml of elution buffer (50 mM Tris-Cl, 500 mM NaCl, 5% glycerol, 400 mM Imidazole, 5 mM TCEP). The eluted protein was then diluted with low-salt buffer (50 mM Tris-Cl, 100 mM NaCl, 5% glycerol, 5 mM TCEP) to achieve a final salt concentration of 300 mM NaCl. The diluted sample was then loaded onto a 5 ml HiTrap Q HP anion exchange column (Cytiva) using a peristaltic pump. The column, connected to an ӒKTA pure™ 25 chromatography system, was washed and eluted with a gradient of NaCl from 300 mM to 2M in 25 mM Tris-Cl buffer (pH8.0) and 2 mM 1,4-dithiothreitol (DTT). The fractions of the elution peak were combined, concentrated and further purified by size exclusion chromatography (SEC) using a Superose 6 10/300 column in a buffer containing 25 mM Tris-Cl (pH8.0), 250 mM NaCl, and 5 mM DTT. The intact complex was identified based on ultraviolet absorbance values at 260 nm and 280 nm, as well as SDS-PAGE analysis. The fraction with highest protein concentration was used immediately for cryo-EM grid preparation. All remaining fractions were snap-frozen in liquid nitrogen and stored at −80 °C.

The protein purification of the PtuA and PtuB complex was carried out similarly as described above. To avoid the toxicity of co-expressing PtuA and PtuB, we expressed the two proteins separately and the cells were combined for co-purification. The wild-type PtuAB proteins could also be expressed when tRNA^Tyr^ is co-expressed. For the non-toxic PtuAB mutants, cells co-expressing both proteins were harvested as described. After HisTrap affinity purification, the eluted protein was loaded to a 5 ml HiTrap Heparin column instead of the HiTrap Q column. All other steps remained unchanged.

### Cryo-EM sample preparation, data collection, and 3D reconstruction

Cryo-EM grids were prepared using a Mark IV Vitrobot. 3 μl of the sample (3 mg/ml) were applied to R 1.2/1.3 Carbon Quantfoil grids, which had been freshly glow-discharged in a Gatan Solarus 950. The grids were double-sided blotted for 4s with a constant force of 0, in a chamber maintained at 100% relative humidity at 8 °C. They were then plunged into liquid ethane and stored in liquid nitrogen before screening.

Grids were initially screened on an Arctica Talos microscope equipped with a K3 camera operated at 200 kV. Good grids were subsequently transferred to a Titan Krios microscope, also equipped with a K3 camera and operated at 300 kV. Micrographs were automatically collected using SerialEM in low-dose mode at a nominal magnification of 130,000× in a super-resolution mode, with an energy filter of 15 eV, leading to a corrected physical pixel size of 0.828 Å/pixel. A total dose of 50 e^-^/ Å^2^ was applied over 50 frames, with a random defocus range of −1.0 to −1.8 µm.

The raw micrographs, initially in super-resolution, were frame-aligned using Patch Motion Correction with cropping to half the original resolution (0.828 Å/pixel), followed by Patch CTF Estimation. Particle picking was performed automatically using Blob Picker with a particle range size set to 100 −180 Å. Subsequent particle inspection and extraction yielded a total of 639,730 particles. The extracted particles underwent two-rounds of 2D classification, resulting in 50,913 high-quality particles. These refined 2D templates were then used for particle picking via Template Picker, followed by another round of particle inspection, extraction, and two-rounds of 2D classification. Ultimately, 199,577 good particles were selected for 3D reconstruction using Ab-initio reconstruction, Homogeneous Refinement, and Non-uniform Refinement. The resulting volume and selected particles exported to Relion for further processing of the complete datasets, where they served as initial model and templates for topaz training.

The complete set of 20,372 raw movies of the Ec78 complex was imported to Relion (v5.0)^31^ for processing. The raw micrographs, initially in super-resolution, were frame-aligned using MotionCor2 (v1.6.4)^32^ with a binning factor of 2. CTF estimation was then performed using Gctf (v1.18)^33^ within Relion (v4.0). Based on a CTF resolution cutoff of 6.0 Å, a total of 19,745 micrographs were selected for further analysis. A subset of 999 randomly selected micrographs was used for topaz training. For topaz training, the cryoSPARC^34^ particle file was first converted to Relion-compatible star file using the PyEM package (csparc2star)^35^ before being imported to Relion. Subsequent 2D classification provided high-quality templates for automated particle picking from the selected 999 micrographs. The picked particles were inspected and refined through another round of 2D classification, and the resulting high-quality particles were used as input for Topaz training. The trained topaz model was then applied for automated particle picking across the entire dataset, yielding a total of 3,630,467 particles. The picked particles were extracted and downscaled by a factor of 2, resulting in a final pixel size of 1.656 Å/pixel. The extracted particles underwent three-rounds of 2D classification, yielding a total of 1,235,478 high-quality particles for further processing. 3D classification was preformed using the initial volume from cryoSPARC, applying alignment and Blush regularization. Similar maps were combined, and the junk particles were discarded, resulting in two good classes: one consisting of 875,922 particles and the other 294,359 particles. For high-resolution reconstruction, those particles were separately re-extracted at the original physical pixel size of 0.828 Å/pixel. Subsequent 3D refinement and post-processing of these two classes generated two maps, leading to final resolution of 2.75 Å and 2.69 Å, respectively, following Non-uniform Refinement.

A data set of 17040 raw micrographs was collected for the PtuAB complex. The same data processing and refinement were carried out, leading to a major class of particles with a reported resolution of 3.25 Å.

Both complexes were modeled based on the Septu PtuAB complex coordinates (8EE7) and the Ec86 coordinates (7V9X), manually adjusted in COOT^36^ and refined in PHENIX^37^.

### Phylogenetic analysis

The phylogenetic analysis was performed with an in-house snakemake pipeline modified from https://github.com/vihoikka/Cas10_prober. Briefly, 91,902 annotated and marked as ‘complete’ genomes were downloaded on August 22, 2024. The genomes were analyzed through Defensefinder (ref) to identify RT and PtuAB sequences. For retron RT analysis, all organisms that annotated as retron systems were singled out, resulting in 8,143 sequences. These RT sequences were clustered with CD-hit (similarity cutoff 0.99) resulting in 1,935 representative sequences. To create a RT phylogenetic tree, proteins were aligned using Muscle 5.1 with the Super5 algorithm intended for large data sets. The RT tree was constructed from the alignment using FastTree 2.1.11 with the WAG+ CA model and Gamma20-based likelihood and visualized in R4.1.1 using ggtree and ggplot.

PtuA and PtuB phylogenetic analyses were carried out similarly. A total of 6295 PtuA and PtuB sequences from either Septu (5878) or Type I-A retrons (417) were extracted and clustered before the phylogenetic tree construction as description above.

### ATPase assay

ATPase activity was measured using the ADP-Glo Kinase assay (Promega). The reactions were carried out with or without 500 μM ultrapure ATP in a reaction buffer containing 20 mM Tris pH8.0, 150 mM NaCl, 2 mM DTT, 10 mM MgCl_2_. Reactions containing 0.2 mg/ml of a protein complex were incubated at 37 °C for 45 minutes before adding ADP-Glo reagent to remove free ATP, followed by the detection reagent. Luminescence was measured on Bio-tek multiplate reader. The reported luminescence reading for each sample was normalized against the readings from reactions without ATP. Mean values obtained from three technical replicates with bars indicating standard deviations.

### Cell growth spot assay

An individual colony from previously transformed bacteria grown on agar plates with appropriate antibiotic(s) was inoculated in LB medium containing the appropriate antibiotic(s) and allowed to grow overnight at 37 °C with shaking. The next day, the overnight culture was used to inoculate a fresh culture that grew until the OD600 reached 0.25. Tenfold serial dilutions of these cultures were prepared in LB medium, and 7.5 μl of these dilutions was spotted onto LB agar plates containing the appropriate antibiotic(s) and inducer (IPTG). The plates were incubated overnight at 37 °C and were imaged the next day using a ChemiDoc Imaging System (Bio-Rad).

For tRNA rescue experiments, plasmids encoding E coli tRNAs were constructed as described in Masuda et al.^21^ and co-transformed with that encoding PtuAB.

### Non-coding RNA and msDNA analysis

The nucleic acids present in either RT or the intact retron-expressing cells were extracted using a Qiagen Plasmid Plus Mid kit and eluted. The nucleic acids co-purified with either RT or the intact retron were extracted by denaturing the purified proteins. The resulting nucleic acids were resolved on a 15% urea PAGE gel and visualized by SYBR Gold staining (ThermoFisher).

### tRNA cleavage

Purified PtuAB, PtuB, and Ec78 at 1 mM concentration were incubated with either the native E coli tRNA^Tyr21^ to test their activity on tRNA cleavage. In vitro transcribed or the native tRNA^Tyr^ was heated to 95°C for 5 minutes, and then immediately transferred to 65°C. After adding MgCl_2_ to a final concentration of 10 mM, the tRNA was allowed to cool to room temperature. The cleavage assay was performed in a buffer containing 10 mM HEPES (pH 7.5), 50 mM KCl, 10 mM MgCl_2_, 1mM DTT and 10 μM MnCl_2_ and at 37 °C for 1 hour. To test the effect of ATP on tRNA cleavage, 1 mM ATP was added. The reaction was stopped by the addition of RNA loading dye (Thermofisher) and heat-denatured at 95 °C for 3 minutes. The reaction products were resolved on 15% urea-polyacrylamide gel electrophoresis gels and visualized by SYBR Gold staining and imaged using the Bio-Rad ChemiDoc system.

### Plasmid construction for human cell transfection

Plasmids used for human cell transfection were adapted from the design described by Khan et al.^10^, in which the reverse transcriptase (RT) is expressed under a CAG promoter, while the non-coding RNA (ncRNA) and Cas9 guide RNA (gRNA) are driven by U6 and H1 promoters, respectively. The RT gene strand was codon-optimized and synthesized with the associated ncRNA through Twist Bioscience. The a1/a2 region of the ncRNA was extended to 18 bp, and the msd donor stem was shortened to 9 bp, following modifications reported in the previous study^10^. The best-performing retron from Khan et al. ^10^ (1531) was used as a performance benchmark. Both tested retrons carried ncRNA sequences designed to introduce a precise three-base edit in the EMX1 locus, converting GAAGGG to AAAGTT.

### Human cell transfections

HEK293T cells constitutively expressing SpCas9 from a CBh promoter at the AAVS1 Safe Harbor locus (GeneCopoeia, SL502) were maintained in high-glucose DMEM supplemented with GlutaMax (Thermo Fisher Scientific, 10569010). For retron transfections, 200,000 cells per sample were nucleofected with 500 ng of plasmid using a Lonza 4D-Nucleofector™ X Unit (AAF-1003X) with the SF Cell Line Kit and pulse code CM130.

For DNA repair inhibition experiments, AZD7648 (1 μM; Selleck Chemicals, S8843) and PolQi2 (3 μM; MedChem Express, HY-150279) were added to the culture media before seeding nucleofected cells into 24-well plates. After 48 hours, genomic DNA (gDNA) was extracted using the Quick-DNA Microprep Kit (Zymo Research) and eluted in 30 μl of ultra-pure, nuclease-free water.

A two-step PCR was performed to amplify the target region and incorporate adapters for Illumina sequencing. Amplicons were sequenced on an Illumina NovaSeq platform using paired-end sequencing (150 bp read length) at a depth of 1.1–3.5 million reads per sample. Sequencing data were analyzed by an in-house Python script to quantify the percentage of precise editing.

## Author contributions

B.W. R.H. and H.L. designed the experiments, B.W. expressed and purified the proteins and performed all biochemical and cryoEM studies; R.H. designed and performed human genome editing experiments; Y.H. provided the native tRNA and tRNA-expression plasmids. B.W. R.H. and H.L. wrote the manuscript. All authors edited the manuscript and provided insightful comments.

## Acknowledgments

Cryo-EM data were collected at the David Van Andel Advanced Cryo-Electron Microscopy Suite (RRID:SCR_023210) at the Van Andel Institute (Grand Rapids, MI). We thank the Van Andel Institute Cryo-EM Core staff for their assistance with data collection. Authors thank all lab members for their support and discussions.

## Data availability

The atomic coordinates of the cryo-EM structures of Ec78 and PtuAB have been deposited in the Protein Data Bank under the identifiers 9NNB, 9NNH, 9NNK and in the Electron Microscopy Data Bank under the entries EMD-49570, EMD-49575, EMD-49576 respectively.

